# GENE-FAM: An automated pipeline for mining gene families and its application to MADS-box genes in *Cannabis sativa*

**DOI:** 10.64898/2026.06.10.731441

**Authors:** Louise Ryan, Nina Trubanová, Grace Pender, Rainer Melzer, Graham M Hughes, Susanne Schilling

**Affiliations:** School of Biology and Environmental Science, University College Dublin, Ireland; Science for Life Laboratory (SciLifeLab), Department of Molecular Biosciences, The Wenner-Gren Institute, Stockholm University, 106 91 Stockholm, Sweden; UCD Earth Institute, University College Dublin, Ireland

**Keywords:** GENE-FAM, gene families, gene annotation, MADS, transcription factors, *Cannabis sativa*

## Abstract

Understanding how gene families evolve can offer great insight into adaptation at the phenotypic and ecological levels. This is particularly true in plants, where transcription factor gene families are often targeted for breeding programs to improve the agronomic traits of economically important crops. While recent advances in next generation sequencing have accelerated the wealth of genomics data, there remains a lack of accessible and reproducible genome mining pipelines tailored for gene family characterisation.

Here, we address this gap by developing GENE-FAM, an automated, scalable and open-source pipeline designed to mine and predict gene families based on conserved domains and motifs. To illustrate its application, we apply GENE-FAM to annotate MADS-box transcription factor genes across multiple *Cannabis sativa* genomes.

A comprehensive set of MADS-box genes was identified across three *C. sativa* cultivars, including both previously annotated and newly predicted genes. Through phylogenetic analyses, we confirm that all type II MADS-box gene subfamilies represented in flowering plants are present in *C. sativa*. Comparing our annotations with those of *Arabidopsis thaliana* and *Solanum lycopersicum* revealed that while most MADS type II families are highly conserved, *SEPALLATA*-like genes have undergone diversification in *C. sativa*. Together, these results demonstrate the application of GENE-FAM for genome-wide identification and characterisation of gene families in non-model species, revealing novel insights into MADS-box gene family evolution in *C. sativa*.

## Introduction

Recent advances in next generation sequencing have led to an exponential increase in the availability of publicly accessible genomic datasets^1^. In order to effectively harness such data, the development of automated and scalable bioinformatics pipelines has become essential. Genome mining, the process of identifying and annotating targeted genes of interest, represents one key area of development in this context, allowing researchers to identify, compare, and categorise genes across species^2–5^. One important application of this approach is gene family characterisation, where the evolution of functionally or structurally similar gene members is assessed^6–8^. By investigating sequence diversity across gene family members, and identifying lineage specific gene duplication and loss events, insights into genetic adaptation and gene function can be inferred^9–11^. As such, gene family analysis is a powerful approach towards identifying potential targets for further experimental validation.

Transcription factor-encoding gene families are of particular interest in the field of plant genetics, as they are essential regulators of plant development, making them valuable targets for crop improvement and speed breeding programmes. MADS-box transcription factors are one well characterised example, with roles in gene regulation across a wide range of developmental pathways from germination to flower and seed development as well as stress responses^12,13^. MADS-box transcription factor genes have been widely characterised across diverse plant species^9,12,14,15^, however few studies have explored their role in *Cannabis sativa*^16–19^. *C. sativa* is a highly versatile crop with numerous industrial and medicinal applications, mainly of cannabinoids and other secondary compounds found in trichomes most abundant on female *C. sativa* flowers^20–23^. Many agronomic traits of *C. sativa* are influenced by flowering and developmental timing^20,24^, hence, MADS-box gene classification is of particular relevance in this species.

Despite the wealth of research characterising gene families across plant species, many studies still implement a manual approach to gene family mining and classification^25–31^, hampering scalability and reproducibility across the field. Moreover, these studies often rely on existing genome annotations, limiting the range of species to those available on public databases such as RefSeq^32^ or Ensembl^33^. Traditionally, studies used keyword-based searches for gene identification in annotated genomes, however these approaches can be unreliable as gene nomenclature systems can be inconsistent across sources^34^. Homology search methods have now become the gold standard approach for genome mining. Additional issues can also arise if genome assembly and annotation quality are not considered, resulting in false reporting of gene duplication and loss events across lineages^34^. As such, there is a clear need for standardisation of methods, particularly with the goal of improving reproducibility and reliability of results across the field.

Here, we address these limitations by developing GENE-FAM, an accessible, fully automated and scalable pipeline tailor-made for gene family characterisation. We apply GENE-FAM to facilitate the genome-wide identification of MADS-box transcription factor genes across three *C. sativa* cultivars. Using a phylogenetics approach, we further classify these genes at the subfamily and member levels, revealing previously uncharacterised patterns of MADS-box gene evolution in *C. sativa*.

## Methods

### The GENE-FAM pipeline

GENE-FAM (Figure 1) was developed to mine gene families based on conserved domains and motifs. The pipeline implements a series of homology searches and gene prediction steps to detect both annotated and previously unannotated loci across target genome assemblies. Each stage of the pipeline is described in detail below.

**Figure 1.**
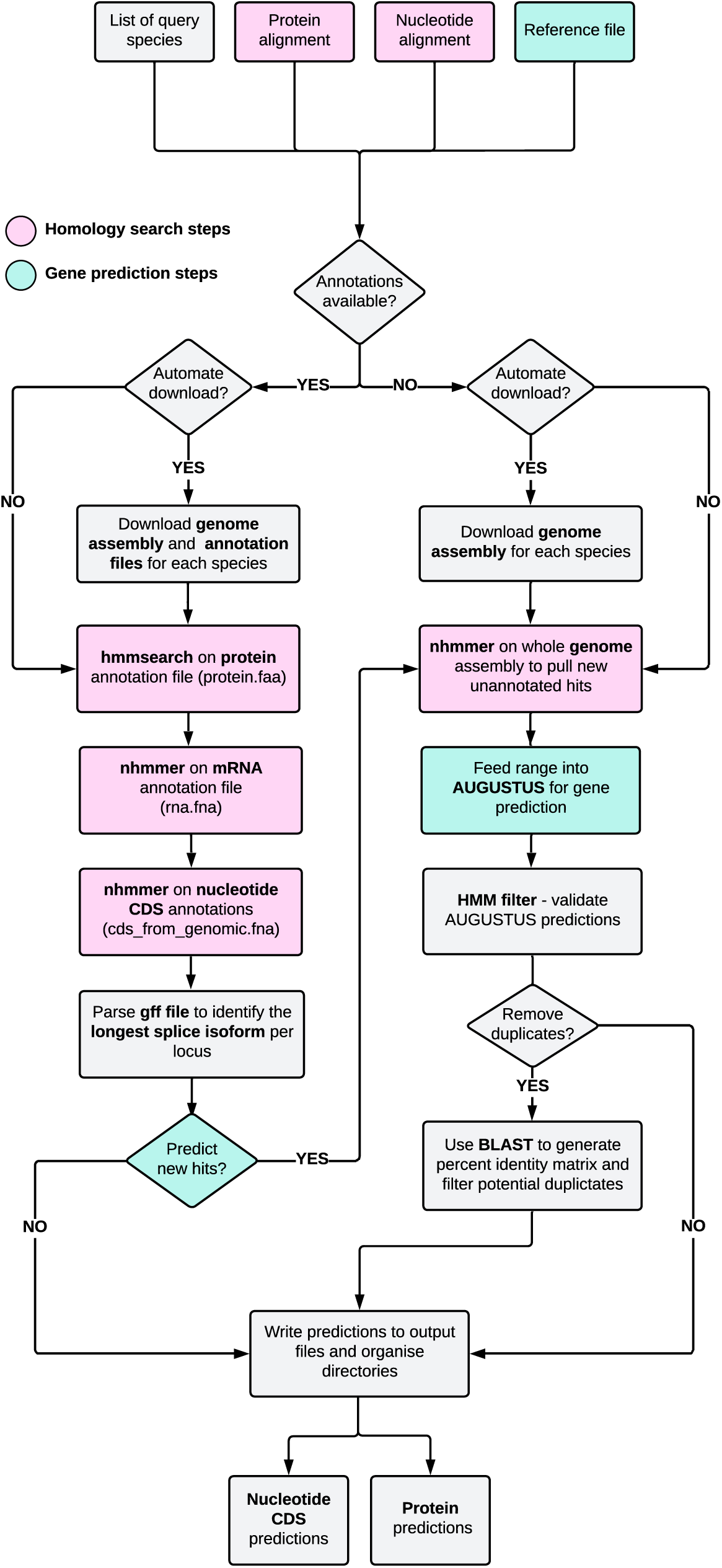
Flowchart illustrating the steps implemented in the GENE-FAM pipeline. Steps and input files highlighted in pink are associated with homology searches, while those in turquoise are associated with gene prediction.

### Mining RefSeq annotations with HMMER

GENE-FAM implements HMMER^35^ to generate profile Hidden Markov Models (HMMs) from two core input files, 1) a protein alignment and 2) a nucleotide alignment of the conserved domain/motif of choice. For a given query species, multiple HMM searches may be performed depending on the availability of existing gene annotations. If the query genome is annotated on NCBI^36^ (www.ncbi.nlm.nih.gov/home/genomes/), these annotations are mined directly, prior to scanning the genome for novel loci. In this case, four additional input files are required including the protein, mRNA, nucleotide coding sequence (CDS), and GFF annotation files. These files, along with the corresponding genome assemblies, can be obtained from the RefSeq database^32^ or automatically downloaded using GENE-FAM for a list of query species. HMM searches are performed on each annotation file. Significant hits surpassing a modifiable e-value threshold (1 x 10^-5^ by default) are compared to retrieve a dataset of unique loci. Using the GFF annotation file, splice isoforms are grouped per locus and the longest isoform is retained. If no RefSeq annotations are available for a given species, GENE-FAM will bypass these annotation search steps and directly scan the assembly for target loci.

### Mining genome assemblies for novel, unannotated hits

To identify novel loci, the nucleotide profile HMM is used to scan the entire genome assembly. Significant hits are retrieved, and coordinates are used to identify target loci within the genome. By default, a range of 5,000 bp upstream and 20,000 bp downstream is retrieved by extending from the 5′ and 3′ coordinates, respectively. These values are provided as a guide and should be adjusted according to the gene structure and target species to ensure the entire gene is captured. Each identified sequence range is iteratively fed into AUGUSTUS^37^ to predict the nucleotide CDS and corresponding amino acid sequence for each novel locus.

### Gene prediction with AUGUSTUS

To guide AUGUSTUS^37^ gene predictions, a customised reference file containing homologous coding sequences from the gene family of interest must be provided as an additional input. This reference file may contain sequences from multiple species, given that they belong to the target gene family. AUGUSTUS^37^ uses this reference file to generate hints, aiding in the identification of intron-exon junctions. The entire reference file may be used to generate hints, or the user may restrict inclusion to the top-scoring ‘n’ hits (‘all’ by default) or to hits above a specified identity threshold (60% by default). BLAT^38^ is used to generate these identity scores based on local alignments between each reference and query sequence. Annotated sequences mined in prior HMM searches may also be included to guide gene prediction and optionally appended to the reference file using a built-in GENE-FAM feature. AUGUSTUS^37^ gene models are trained on various species. Users should select the training species most closely related to their species of interest to improve accuracy of gene predictions.

### Validating the specificity of predictions

Each AUGUSTUS^37^ prediction is scanned to ensure the motif identified in the HMMER^35^ search is present in the final output. By default, 90% of the motif region must be included to be considered for downstream analyses. Additionally, each AUGUSTUS^37^ prediction undergoes a final HMM filter step, where HMMER^35^ is used to validate the presence of the motif/domain in the prediction. Hits that surpass the specified e-value threshold are considered valid. All mined sequences can be classified as ‘functional’ or ‘pseudogenised’ based on the presence of in-frame stop codons and sequence length. A user-modifiable default length threshold of 300 bp is used to identify possible pseudogenes in predicted sequences.

### Filtering out potential duplicates arising from assembly error

GENE-FAM has an optional feature to remove duplicates which may arise from assembly errors (e.g. misassembled alleles or multiple gene copies due to the presence of redundant contigs). If selected, GENE-FAM uses BLASTP^39^ to generate an all-by-all identity matrix, calculating amino acid identity scores for all pairwise comparisons between predictions. By default, an amino acid identity threshold of ≥90% is applied when identifying duplicates, however, this can be customised. Two methods are available to identify duplicates, including the “pairwise” and “clustered” algorithms (Figure S1). In the pairwise approach, duplicate pairs are identified as mutual best scoring hits in the identity matrix. More than two members may exist in a given “pair” if each member shares the same maximum identity score. In the clustered algorithm, genes with sequence identity values above the user-defined threshold are grouped into “clusters”. In each pair or cluster, the member located on the longest contig is retained to prioritise chromosomal loci over those on unplaced scaffolds.

### Generating GENE-FAM input files for MADS-box gene mining

To apply GENE-FAM to mine MADS-box transcription factors across *C. sativa* cultivars, nucleotide and protein alignments of the MADS domain were required. A custom nucleotide alignment was generated using 108 *Arabidopsis thaliana* (thale cress) MADS-domain coding sequences, retrieved from the NCBI RefSeq database^32^. These sequences were aligned using MAFFT, v7.562 ^40–42^, applying the E-INS-i iterative refinement method. Sequences were manually cropped in Jalview, v2.11.3.2^43^ to exclusively retain the MADS-domain region. The input protein alignment for the ‘SRF-type transcription factor’ domain (PF00319) was downloaded directly from the InterPro database^44^. To generate the reference file used to guide AUGUSTUS gene prediction, annotated MADS-box coding sequences from *A. thaliana* (TAIR10.1; GCF_000001735.4)*, S. lycopersicum* (tomato; SL3.1; GCF_000188115.5)*, Malus domestica* (apple; ASM211411v1; GCF_002114115.1), and *C. sativa* were downloaded from NCBI^32,36^. Annotations from the *C. sativa* CBDRx genome (GCF_900626175.1) were included to improve gene prediction accuracy across unannotated genomes from other *Cannabis* cultivars.

### Running GENE-FAM on A. thaliana, S. lycopersicum and C. sativa genomes

Reference genome assemblies and associated annotation files were downloaded from the NCBI RefSeq database^32^ for *A. thaliana*, *S. lycopersicum*, and *C. sativa* (Table 1). While the *C. sativa* and *S. lycopersicum* reference genomes used in this analysis have since been superseded by newer assemblies, the versions we used represented the current reference genomes at the time of analysis and enable direct comparison of our results to a broader body of published research. *A. thaliana* and *S. lycopersicum* were selected as test species to benchmark pipeline performance due to their well annotated genomes. To assess pipeline performance on unannotated assemblies, four additional genomes from two alternative *C. sativa* cultivars were selected (Table 1). Haplotype resolved assemblies of ‘Kompolti’ (KOMPa and KOMPb) and ‘White Widow’ (WHWa and WHWb), were retrieved from the literature^45^.

**Table 1.** Summary of genome assemblies analysed with GENE-FAM in this study. Species and common names are provided, along with genome assembly names and accession identifiers. NCBI BioProject identifiers are provided where assembly identifiers are not available. Genome annotation status is indicated for each species. As multiple *C. sativa* genomes were included, cultivar names are also provided.

| Species Name | Common Name | Cultivar | Assembly Name | Assembly/<br>BioProject ID | Annotation<br>Available | Genome Source |
| --- | --- | --- | --- | --- | --- | --- |
| *Arabidopsis thaliana* | Thale cress | NA | TAIR10 | GCF_000001735.4 | Yes | NCBI RefSeq<sup>32</sup> |
| *Solanum lycopersicum* | Tomato | NA | SL3 | GCF_000188115.5 | Yes | NCBI RefSeq<sup>32</sup> |
| *Cannabis sativa* | Cannabis | CBDRx | cs10 | GCF_900626175.2 | Yes | NCBI RefSeq<sup>32</sup> |
| *Cannabis sativa* | Cannabis | Kompolti | KOMP<sub>a</sub> | PRJNA992904 | No | Lynch *et al.*, 2025 |
| *Cannabis sativa* | Cannabis | Kompolti | KOMP<sub>b</sub> | PRJNA992905 | No | Lynch *et al.*, 2025 |
| *Cannabis sativa* | Cannabis | White Widow | WHW<sub>a</sub> | PRJNA1140642 | No | Lynch *et al.*, 2025 |
| *Cannabis sativa* | Cannabis | White Widow | WHW<sub>b</sub> | PRJNA1140642 | No | Lynch *et al.*, 2025 |

GENE-FAM was run on each assembly using default parameters and thresholds, with A. thaliana selected as the AUGUSTUS trained species. RefSeq annotation files were mined where available (A. thaliana, S. lycopersicum, and C. sativa CBDRx). Potential duplications arising from assembly error were flagged using the GENE-FAM ‘clustered’ algorithm for duplicate detection with a threshold of ≥90% sequence identity (Supplementary Figure 1). A known HMMER error was encountered when parsing the KOMPb genome due to a lack of guanine residues in the first 6,586 bp of Chromosome 1. This was resolved by concatenating a short pseudo-contig (‘ATGCATGCATGCATGC’) to the beginning of the assembly. This addition may have slightly impacted HMMER^35^ e-values, though the effect is expected to be negligible. The chromosomal distribution of identified MADS-box genes across the *C. sativa* CBDRx genome was visualised using MG2C, v2.1^46^ (Figure 2). Overlapping gene sets between *C. sativa* cultivars were visualised using the InteractiVenn^47^ resource.

**Figure 2.**
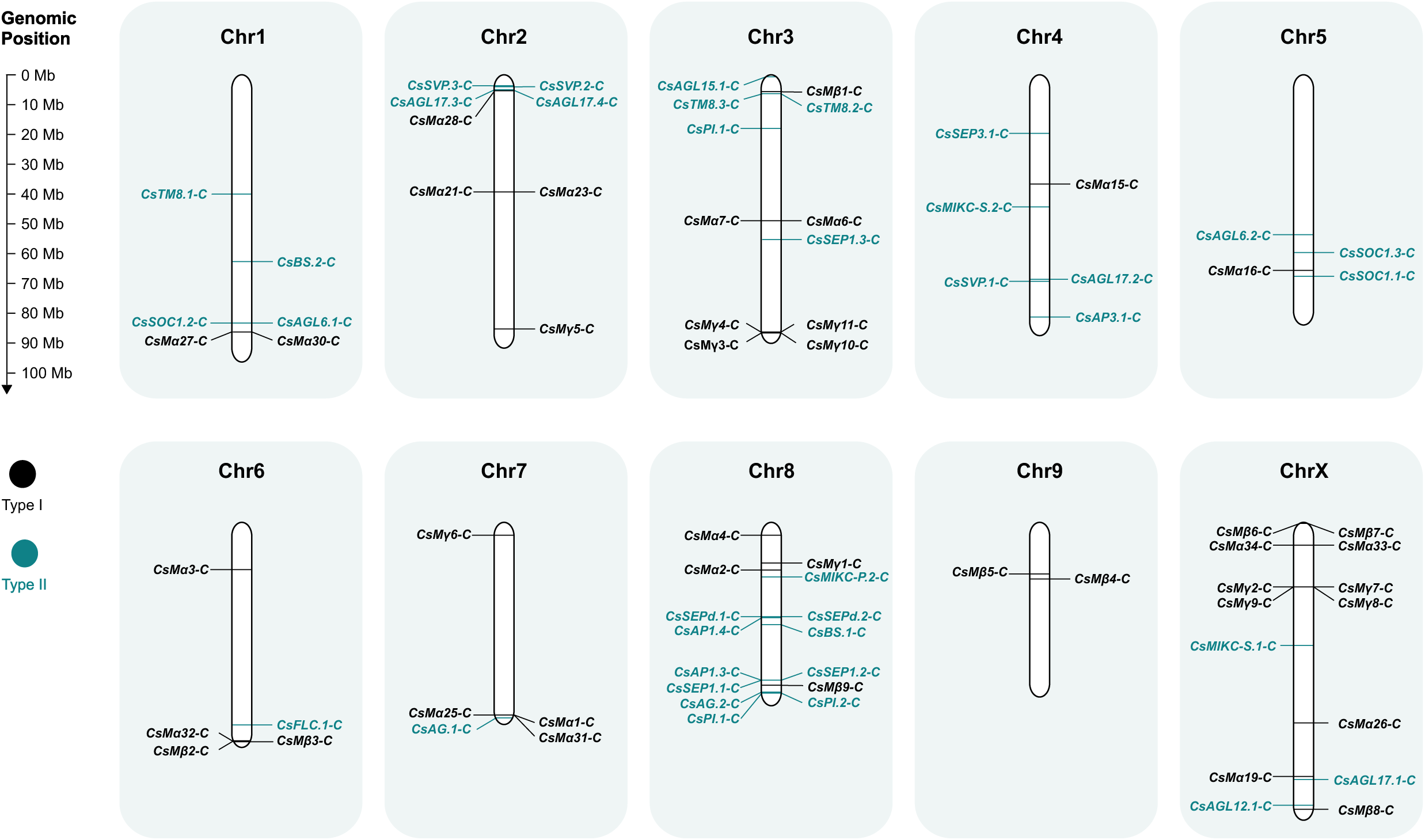
Chromosomal distribution of MADS-box genes across the *C. sativa* CBDRx genome. Type I and type II MADS-box genes are labelled in black and turquoise respectively. *C. sativa* CBDRx genes *CsMα.15-C*, *CsAP1.1-C*, and *CsAP1.2-C* are not included as these are located on unplaced scaffolds. Chromosome lengths are displayed to scale, and a genomic position map is provided for reference.

### Sequence alignment and phylogenetic analysis

Output amino acid predictions mined with GENE-FAM were aligned using MAFFT^40–42^ (E-INS-i iterative refinement method), including only the longest isoform per locus. Each alignment was manually trimmed in Jalview^43^ to retain only the 60 aa MADS domain region^48^. Each alignment was masked using AliStat v1.3^49^ with completeness scores for individual sites (C_c_scores) determined for each alignment (Tables S2-S5). Maximum likelihood gene trees were then constructed from these alignments using IQ-TREE^50–52^. Branch support was assessed using both ultrafast bootstrap approximation and the Shimodaira-Hasegawa approximate likelihood ratio test (SH-aLRT), each with 1,000 replicates. The resulting trees were visualised using iTOL^53^ (Figures S2, S3). Phylogenetically derived classifications for type II MADS-domain proteins were further validated by confirming the presence of the M- and K-superfamily domains across *A. thaliana, S. lycopersicum,* and *C. sativa* using the NCBI Batch CD-Search tool^54–56^ (Table S1).

### Assigning informative gene names to identified MADS-box genes

An informative nomenclature system was derived to classify each identified MADS-box gene across species and *C. sativa* cultivars. Using this system, genes are assigned based on four components: (1) an abbreviated species identifier, (2) the phylogenetic subfamily name, (3) a numeric identifier to distinguish between paralogs in the clade and (4) an abbreviated cultivar identifier where applicable. For example, a gene named ‘*CsSVP.2-C*’, belongs to *C. sativa* (Cs), is a member of the SVP subfamily, represents a paralog in that subfamily (SVP.2), and belongs to the CBDRX (-C) cultivar.

## Results

### GENE-FAM identifies novel MADS-box genes in *A. thaliana* and *S. lycopersicum*

To benchmark performance in well annotated model species, GENE-FAM was applied to the *A. thaliana* and *S. lycopersicum* reference genomes. A total of 114 MADS-box genes were identified in the *A. thaliana* genome. Of these, 113 were existing RefSeq annotations, while one was a novel putative MADS-box gene. Based on conserved domain structure and phylogenetic analysis (Figures S2, S3), 67 of these identified genes were classified as type I and 47 as type II MADS-box genes. RefSeq coding sequence lengths ranged from 330 bp to 1,395 bp, with a median of 748.5 bp across all identified sequences. The newly identified gene, named here as *AtMα.2*, is located in an unannotated region of Chromosome 3 (Figure S4), contains an intact open reading frame, and has a length of 477 bp. Based on phylogenetic analysis, *AtMα.2* belongs to the Mα subclade of type I MADS-box genes (Figure S2).

Similarly, GENE-FAM identified 131 RefSeq annotations and 4 novel putative functional MADS-box genes in the *S. lycopersicum* genome. While 10 additional novel loci were detected, we predict these to be pseudogenes based on truncated sequence length (Table S1). RefSeq coding sequences had similar lengths to those identified in the *A. thaliana* genome. Each of the 4 newly identified predictions was found to belong to the Mα subclade, and are located in previously unannotated regions of the genome (Figures S5A-D). These results support the ability of GENE-FAM to detect novel putative genes and improve on existing annotations, even in well-annotated model species.

### Genome-wide identification of MADS-box genes across *C. sativa* cultivars

Mining the *C. sativa* CBDRx reference genome using GENE-FAM resulted in the identification of 80 MADS-box genes, 68 of which were previously annotated, with 12 representing newly identified genes (Table S1). Overall, 41 type I and 39 type II MADS-box genes were identified. Of the 12 newly identified genes, only three were predicted to be functional based on sequence length (> 300 bp). Each of these three predictions were located on chromosome assembled regions (Figures 6A-C), contained the MADS domain, and were classified as type I MADS domain proteins belonging to the *Mα* clade (Figure S2). These newly predicted sequences ranged from 600 bp to 1,287 bp, consistent with RefSeq annotation sequence lengths. Mapping the chromosomal location of all identified MADS-box genes revealed widespread distribution across the CBDRx genome (Figure 2). Chromosome 8 (NC_044379.1) contained the highest number of MADS-box genes (n = 15), while chromosome 2 (NC_044376.1) had the lowest (n = 2) (Figure 3).

**Figure 3.**
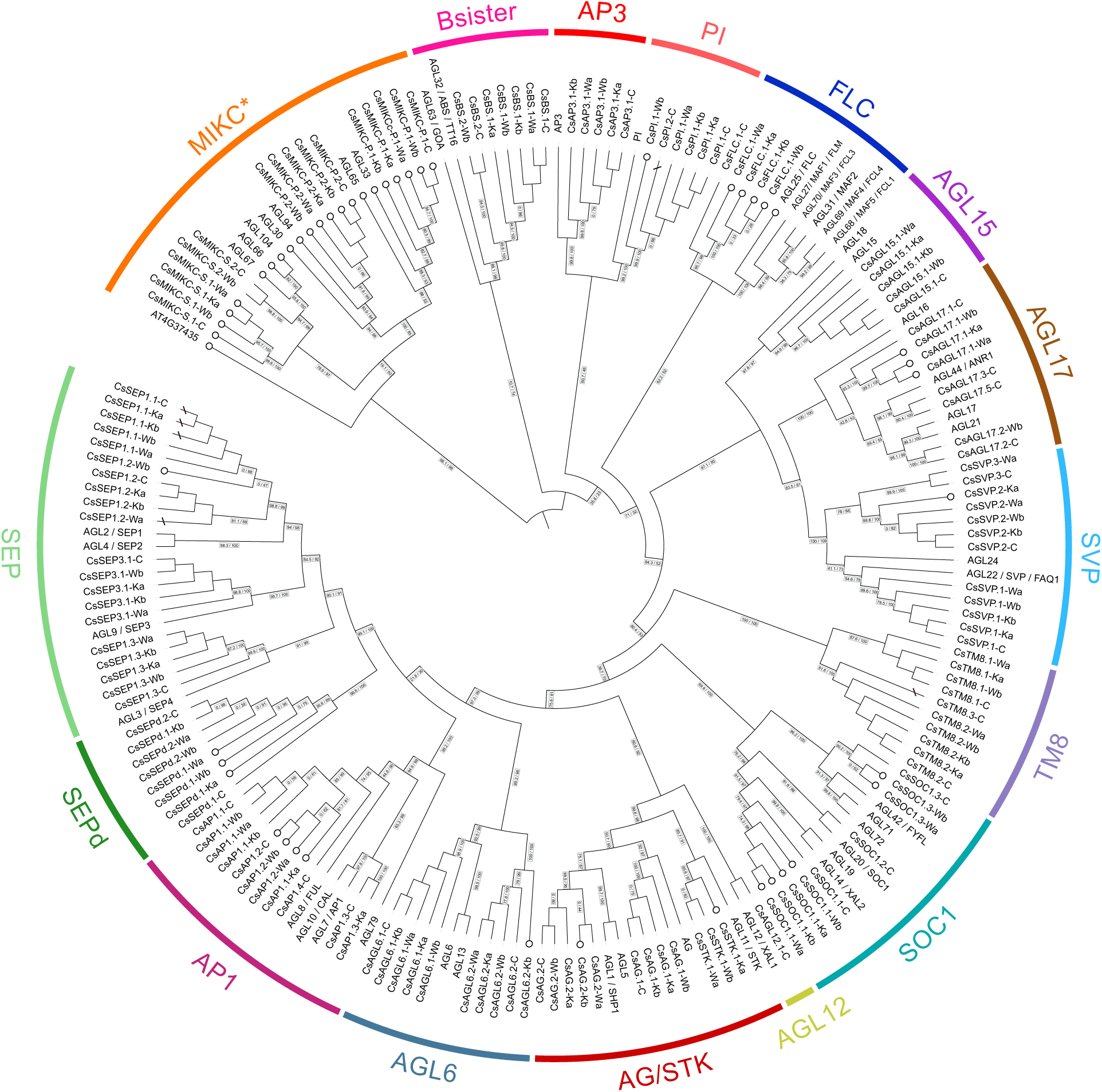
Maximum likelihood phylogeny of Type II MADS-domain proteins from *A. thaliana* and *C. sativa*. Node support values are displayed as SH-aLRT support (%) / ultrafast bootstrap support (%). White circular tip labels next to gene names indicate the absence of at least a partial M- or K-superfamily domains. Diagonal lines across branches represent potential misassembled duplicates flagged using the GENE-FAM ‘clustered’ algorithm based on a threshold of ≥90% sequence identity. *C. sativa* cultivars receive the following abbreviations at the end of each gene name, ‘C’ for the CBDRx genome, ‘Wa’ and ‘Wb’ for the White Widow genomes (WHWa and WHWb), and ‘Ka’ and ‘Kb’ for the Kompolti genomes (KOMPa and KOMPb).

To assess population level variation across different *C. sativa* cultivars, GENE-FAM was applied to mine MADS-box genes in two additional strains, ‘Kompolti’ and ‘White Widow’, using haplotype resolved genomes (KOMPa and KOMPb, and WHWa and WHWb^45^, respectively). Copy number variation was detected across each genome, with 62 sequences recovered from KOMPa, 54 from KOMPb, 67 from WHWa and 62 from WHWb (Table S1). It is difficult to assess whether this variation arises from assembly errors or reflects true structural variation between haplotypes. Variation may also arise from the limitations of *ab initio* gene prediction, or due to genes being fragmented across poorly assembled contigs or scaffolds.

### Phylogenetic classification of MADS-box genes in *C. sativa*

To further categorise and annotate MADS-box genes identified across all *C. sativa* genomes, a series of phylogenetic trees were generated. Using annotations from *A. thaliana* and *S. lycopersicum* to guide classifications, sequences were grouped as either type I (Figure S2) or type II (Figure S3) MADS-box genes. By comparing our results to previous studies^57,58^ we determined that all major subfamilies of type II MADS-domain proteins known in flowering plants were also present in *C. sativa* (Figure 3, Table S1). Including *S. lycopersicum* in our analyses confirmed the presence of the *TM8* MIKC^C^ subfamily in *C. sativa*, which is absent in *A. thaliana* (Figure 3, Figure S3).

Compared to *A. thaliana*, the MIKC* clade was found to be less diversified in *C. sativa* with two members in the P and one in the S subclade (Figure 3). Similarly, in the MIKC^C^ clade, only one FLC-like member was identified across each *C. sativa* genome, whereas this subfamily consists of six genes in *A. thaliana* and three in *S. lycopersicum* (Figure 3, Figure S3).

For the *SEP* subfamily, a particularly interesting phylogenetic pattern was revealed. The highly conserved *SEP3*- and *SEP1/2/4*- (also termed *LOFSEP*) clades were reconstituted, with clear *C. sativa* orthologs in both clades. However, in addition to that, *C. sativa* genes were identified that clearly clustered in the *SEP* subfamily but could not reliably be assigned to either the *SEP3* or *SEP1/2/4* clade (Figure 3). Additional phylogenetic reconstructions using only *SEP*-like and closely related genes from *A. thaliana*, tomato and *C. sativa* confirmed this topology (Figure S7). We termed those genes *CsSEPd.1* and *CsSEPd.2* (for SEP-divergent).

Variation was also detected at cultivar level, with putative lineage specific duplication and loss events recorded across *C. sativa* cultivars. While most type II MADS-box genes (n = 25) were conserved across individuals, cultivar specific repertoires were observed (Figure S8). Seven CBDRx-specific genes were identified, including *TM8.3* and *PI.2*. Tandem duplication of the *TM8.2/3* and *PI.1/2* loci was inferred based on chromosomal positioning (Figure 2) and phylogenetic groupings (Figure 3). Notably, these loci were flagged by GENE-FAM as potential redundant entries as the shared sequence identity exceeded 90%, suggesting that they may represent misassembled alleles rather than true CBDRx-specific paralogs. Gene sets were also largely conserved between haplotypes (n = 27, WHW; n = 20, KOMP), though a limited number of haplotype-specific genes were detected (Figure S9). All absent copies from the KOMPb haplotype, except for *AP1.3,* were X-linked (Table S1), consistent with a male (XY) karyotype. No type II genes were detected on the Y chromosome, though 9 type I genes were Y-linked, spanning the Mα, Mβ and Mγ clades (Table S1).

We further identified a number of putative MADS-domain proteins in *C. sativa* with truncated MADS or K domains, while all mined *A. thaliana* MIKC^C^ MADS-domain proteins, with the exception of FLC, were identified as having a complete M- and K-domain (Figure 3). It is possible that this finding is due to the presence of pseudogenes in *C. sativa*. However, the fact that truncated domains were predominantly observed in the two haplotype resolved genomes (‘White Widow’, ‘Kompolti’) suggests that this is a sign of artefacts of the genome assembly used for these *C. sativa* genomes.

## Discussion

Scalable and reproducible methods for gene family characterisation are essential for keeping pace with the growing availability of plant genomes. To address this need, we developed GENE-FAM, an automated and open-source pipeline for genome wide mining of gene families based on conserved domains and motifs. We applied GENE-FAM to annotate MADS-box transcription factor genes across multiple *C. sativa* genomes and found that it accurately recovered known annotations, predicted novel genes, and enabled rapid comparative analyses across lineages, including previously unannotated genomes.

GENE-FAM successfully identified all previously characterised type II MADS-box transcription factors in each tested genome, when compared to previous studies^59–62^, demonstrating that an automated approach can perform comparably to manually curated methods while significantly improving on efficiency. Moreover, we find that GENE-FAM predicts novel, previously unannotated Type I MADS-box genes across all taxa, highlighting the importance of applying *ab initio* gene prediction in gene family characterisation studies, even in model species. Further validation using the NCBI CDD search tool confirmed the presence of the MADS domain in all final output predictions (Table S1), demonstrating the specificity of our pipeline and its ability to exclude off-target hits from resulting annotations.

Applying GENE-FAM across multiple *C. sativa* genomes allowed for orthologues of well characterised MADS-box genes to be identified, providing candidates for future experimental and functional validation. For example, the *TM8* gene plays an important role in floral development in *S. lycopersicum*^63^. Orthologs of *TM8* are also associated with flower and fruit development in other Rosales members including *Cucumis sativus* (cucumber)^64,65^, *Pyrus pyrifolia* (Asian pear)^66^, and *M. domestica*^67^. The presence of two copies of the *TM8* gene across all mined *C. sativa* genomes (excluding KOMPb) suggests a similar genomic mechanism may underpin reproductive development in this species.

Phylogenetic characterisation of type II MADS-box genes across *C. sativa* cultivars revealed a divergent group of *SEP* homologues, forming a sister group to the highly conserved *SEP3* and *SEP1/2/4* clades. Notably, this clade was specific to *C. sativa*, with no corresponding orthologs detected in *A. thaliana* or *S. lycopersicum*. This observation is particularly interesting in light of recent work showing that *CsSEPd.2* influences inflorescence number per branch and seed yield via the ethylene pathway^68^, and that ethylene related genes also contribute to floral organ identity determination and sexual plasticity in *C. sativa*^69^. These overlapping regulatory networks suggest that *SEP* divergent genes may sit at the intersection of flowering regulation and reproductive development, an area that warrants further experimental validation.

Putative lineage specific gene duplication and loss events were detected across *C. sativa* cultivars. Copy number variation across cultivars may be driven by ecological factors^70^ or domestication and breeding history^71^. As such, the variation detected here may be adaptive, providing interesting genetic targets for characterising phenotypic diversity across *C. sativa* cultivars. However, assembly and annotation errors cannot be ruled out as an explanatory factor driving this variation. For example, duplication of the *TM8.2/3* and *PI.1/2* loci detected in the non-haplotype-resolved CBDRx assembly likely represent misassembled alleles rather than true paralogs. Similarly, *AP1* orthologs map to Chromosome 3 in the ‘White Widow’ and ‘Kompolti’ genomes but to an unplaced contig in CBDRx, while *CsTM8.1* is located on Chromosome 1 in CBDRx but on the X chromosome in both ‘White Widow’ and ‘Kompolti’. Whether these discrepancies reflect true chromosomal rearrangements or assembly artefacts remains to be determined.

As genome sequencing enters the era of reference-quality and telomere-to-telomere assemblies, the availability of rapid, high-throughput methods to analyse them becomes a key bottleneck in translating raw sequence data to biological knowledge. GENE-FAM overcomes this bottleneck by enabling rapid, scalable, and reproducible annotation of gene families across species, supporting future large-scale analyses across diverse genomic datasets.

## Data availability

The GENE-FAM source code and associated files are available on GitHub (https://github.com/Louise-Ryan/GENE-FAM). A preconfigured virtual machine for GENE-FAM is also available via Figshare (https://figshare.com/s/790244b8d1ba32c6ff29). Supplementary data and figures are provided.

## Acknowledgements

This work was supported by Research Ireland under the Centre for Research Training in Genomics Data Science grant [18/CRT/6214] (LR), and a UCD Ad Astra Fellowship grant (GMH). NT is funded by the European Union (SciLifeLab PULSE 101177199). Views and opinions expressed are however those of the author(s) only and do not necessarily reflect those of the European Union or the European Research Executive Agency. Neither the European Union nor the granting authority can be held responsible for them. G.P. was supported by a postgraduate fellowship under the IRC EPA Scheme (grant no. GOIPG/2021/1072).

## Author Contributions

Conceptualisation of the study, all authors. Conceptualisation of the GENE-FAM pipeline, L.R. and G.P. Development, implementation, and validation of the GENE-FAM pipeline, L.R. Application of the pipeline to plant datasets, N.T. Data analysis, interpretation and visualisation, L.R. and N.T. Writing, reviewing, and editing the manuscript, all authors. Supervision, R.M., G.M.H. and S.S. All authors have read and approved the final manuscript.

## Conflict of interest

The authors declare no competing interests.

## Supplementary Figures

**Figure S1. Illustrating the ‘Clustered’ and ‘Pairwise’ algorithms for removing potential duplicates from GENE-FAM output predictions.** In the ‘pairwise’ algorithm, duplicate pairs are identified as mutual best scoring hits in the percent identity matrix. Note that more than 2 members may exist in a given pair, if each member shares the same maximum identity score. Mutual best scores are only considered pairs if they exceed a user defined sequence identity threshold. The member in each pair which is located on the longest contig is retained. In the ‘clustered’ algorithm, genes that share percent identity values greater than the user defined threshold are combined into clusters. The member in each cluster which is located on the longest contig is retained.

**Figure S2. Maximum likelihood phylogeny of type I MADS-domain proteins from *Arabidopsis thaliana*, *Solanum lycopersicum*, and three different *Cannabis sativa* cultivars.** Bootstrap values are displayed at each node in the phylogeny. Detailed information about all displayed genes can be found in Table S1.

**Figure S3. Maximum likelihood phylogeny of type II MADS-domain proteins from *Arabidopsis thaliana*, *Solanum lycopersicum*, and three different *Cannabis sativa* cultivars.** Bootstrap values are displayed at each node in the phylogeny. Detailed information about all displayed genes can be found in Table S1.

**Figure S4. Genomic position of a newly predicted MADS-box gene, *AtMα.2*, in *A. thaliana* using GENE-FAM.** *AtMα.2* is located on chromosome 3 (NC_003074.8) with a predicted coding sequence spanning positions 15,590,521-15,591,241 in the 5’ to 3’ direction. Six-frame translation tracks are shown, including start (green) and stop (red) codons with arrows indicating translation on the forward or reverse strand. *AtMα.2* overlaps with an uncharacterised protein (AT3G43686), predicted by NCBI to encode a pseudogene. Notably, *AtMα.2* and AT3G43686 are encoded on opposite strands, indicating that they do not share the same coding sequence, and supporting the classification of *AtMα.2* as a novel, previously unannotated MADS-box gene.

**Figure S5. Genomic position of newly predicted MADS-box genes in *S. lycopersicum* using GENE-FAM.** Novel predictions *SlMα.5* (A), *SlMα.4* (B), *SlMα.1* (C) and *SlMα.2* (D) are represented. Chromosomal locations are provided for each gene. Six-frame translation tracks are shown, including start (green) and stop (red) codons with arrows indicating translation on the forward or reverse strand. No previously annotated genes are found to overlap with any novel MADS-box gene prediction. Aggregated RNA-seq exon coverage tracks are shown. Overlapping RNA-seq coverage provides additional support for *SlMα.5*, indicating active transcription of the locus.

**Figure S6. Genomic position of newly predicted MADS-box genes in *C. sativa* using GENE-FAM.** Novel predictions *CsMα.1-C* (A), *CsMα.32-C* (B) and *CsMα.22-C* (C) are represented. Chromosomal locations are provided for each gene. Six-frame translation tracks are shown, including start (green) and stop (red) codons with arrows indicating translation on the forward or reverse strand. No previously annotated genes are found to overlap with any novel MADS-box gene prediction. Aggregated RNA-seq exon coverage tracks are shown, though support was minimal or absent for most loci.

**Figure S7. Maximum likelihood phylogeny of proteins from the SEPd, SEP3, LOFSEP, AGL6, and AP1 subfamilies from *A. thaliana, S. lycopersicum*, and *C. sativa* (CBDRx cultivar).** High bootstrap values support the grouping of SEP-divergent (SEPd) proteins as a sister clade to the SEP3 and LOFSEP subfamilies.

**Figure S8. Venn diagram illustrating conserved and cultivar specific Type II MADS-box gene repertoires across the CBDRx, ‘White Widow’ and ‘Kompolti’ *C. sativa* cultivars.** Haplotype specific gene sets for ‘White Widow’, and ‘Kompolti’ were combined to represent cultivar level repertoires.

**Figure S9. Venn diagram illustrating conserved and haplotype specific Type II MADS-box gene repertoires across the (A) ‘White Widow’ and (B) ‘Kompolti’ cultivars.** Haplotypes are labelled as WHWa/WHWb and KOMPa/KOMPb, respectively.

## Supplementary Tables

**Table S1. Overview of MADS-domain genes and corresponding protein sequences mined with GENE-FAM.** Information is provided for *Arabidopsis thaliana* TAIR10.1 (GCF_000001735.4), *Solanum lycopersicum* SL3.1 (GCF_000188115.5), *Cannabis sativa* CBDRx (GCF_900626175.1), and the haplotype-resolved ‘White Widow’ (WHWa, WHWb) and ‘Kompolti’ (KOMPa, KOMPb) genomes.

**Table S2. AliStat summary statistics for type I MADS-domain protein alignments corresponding to Figure S2.** Scores are shown for both non-cropped alignments and alignments cropped using a 75% minimum C-score threshold for individual sites.

**Table S3. AliStat summary statistics for type II MADS-domain protein alignments corresponding to Figure S3.** Scores are shown for both non-cropped alignments and alignments cropped using a 75% minimum C-score threshold for individual sites.

**Table S4. AliStat summary statistics for type II MADS-domain protein alignments corresponding to Figure 3.** Scores are shown for both non-cropped alignments and alignments cropped using a 65% minimum C-score threshold for individual sites.

**Table S5. AliStat summary statistics for SEPd, SEP3, LOFSEP, AGL6, and AP1 protein alignments corresponding to Figure S7.** Scores are reported for both non-cropped alignments and alignments cropped using an 80% minimum C-score threshold for individual sites.

## Notes

### Competing Interest Statement

The authors have declared no competing interest.

