## Supplemental Figures for "GENE-FAM: An automated pipeline for mining gene families and its application to MADS-box genes in *Cannabis sativa*"

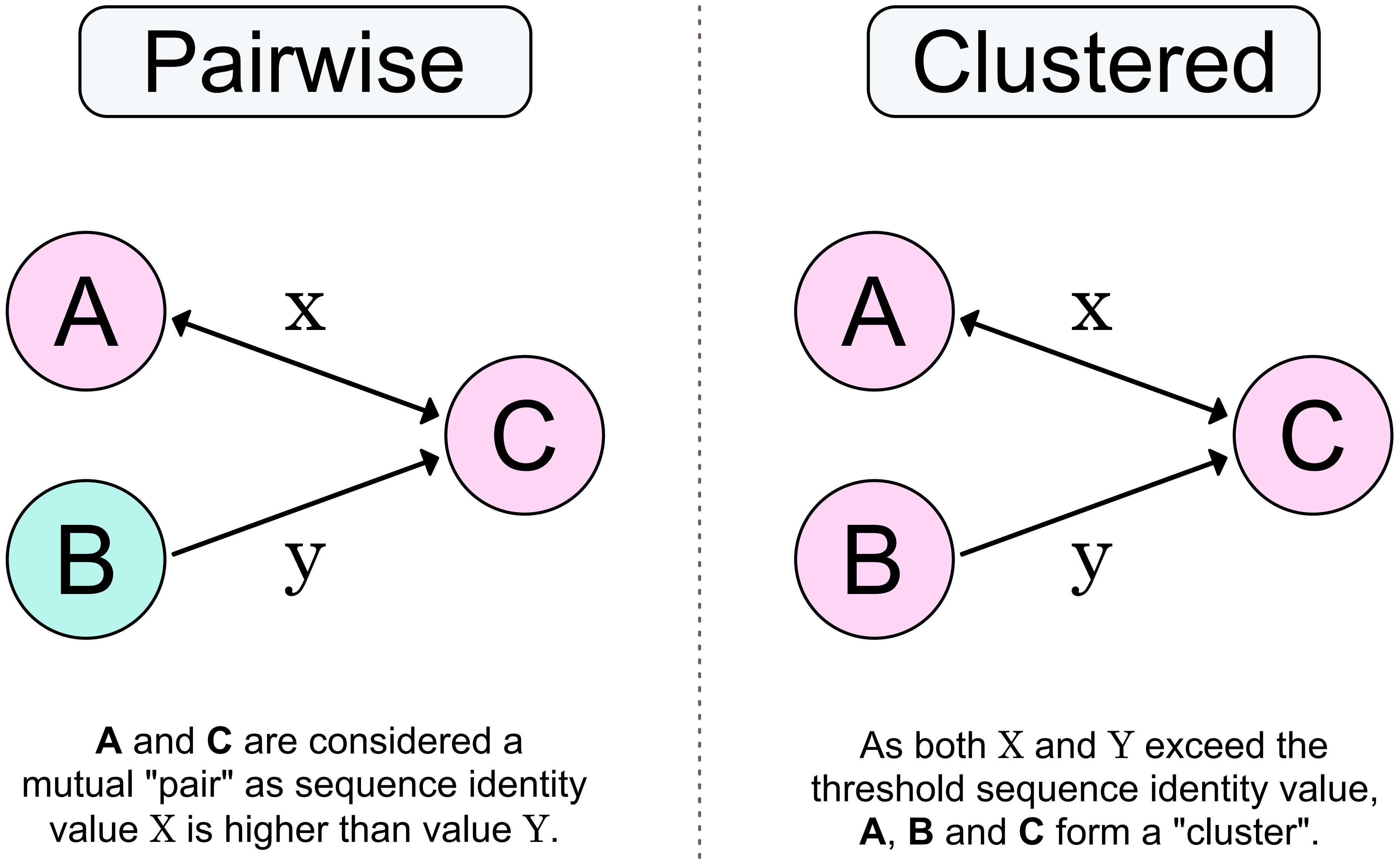


**Figure S1. Illustrating the 'Clustered' and 'Pairwise' algorithms for removing potential duplicates from GENE-FAM output predictions.** In the 'pairwise' algorithm, duplicate pairs are identified as mutual best scoring hits in the percent identity matrix. Note that more than two members may exist in a given pair, if each member shares the same maximum identity score. Mutual best scores are only considered pairs if they exceed a user defined sequence identity threshold. The member in each pair which is located on the longest contig is retained. In the 'clustered' algorithm, genes which share percent identity values greater than the user defined threshold are combined into clusters. The member in each cluster which is located on the longest contig is retained.

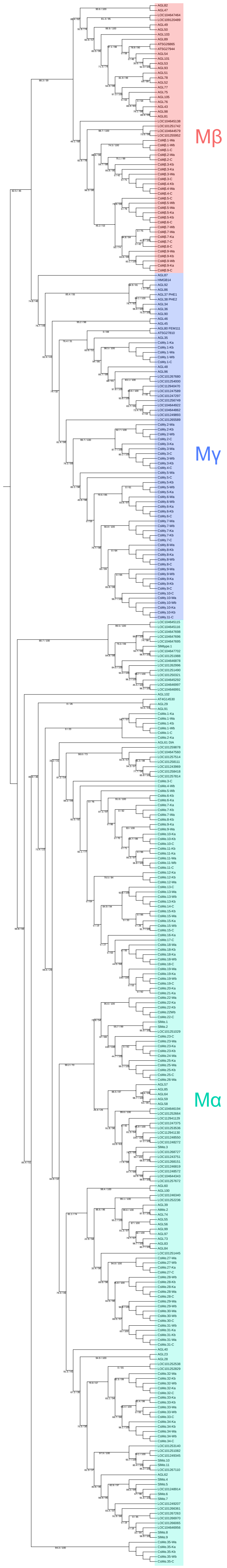


**Figure S2. Maximum likelihood phylogeny of type I MADS-domain proteins from *Arabidopsis thaliana*, *Solanum lycopersicum*, and three different *Cannabis sativa* cultivars.** Bootstrap values are displayed at each node in the phylogeny. Detailed information about all displayed genes can be found in Table S1.

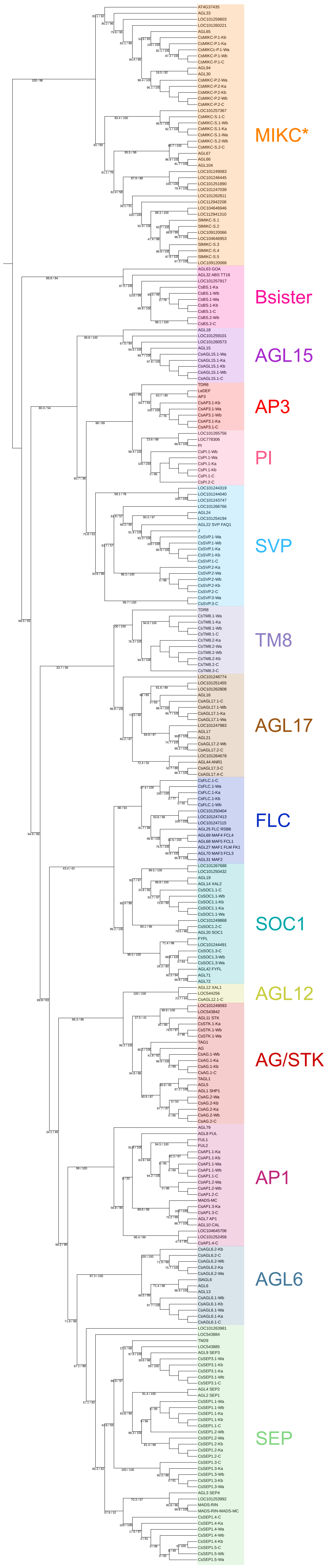


**Figure S3. Maximum likelihood phylogeny of type II MADS-domain proteins from *Arabidopsis thaliana*, *Solanum lycopersicum*, and three different *Cannabis sativa* cultivars.** Bootstrap values are displayed at each node in the phylogeny. Detailed information about all displayed genes can be found in Table S1.

### *AtMα.2,* NC_003074.8:15590521-15591241

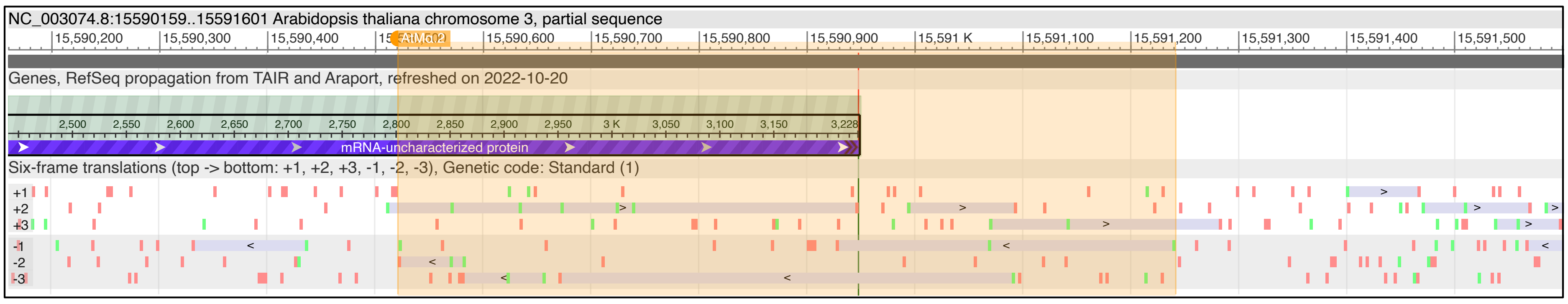


**Figure S4. Genomic position of a newly predicted MADS-box gene, *AtMα.2*, in *A. thaliana* using GENE-FAM.** *AtMα.2* is located on chromosome 3 (NC_003074.8) with a predicted coding sequence spanning positions 15,590,521-15,591,241 in the 5’ to 3’ direction. Six-frame translation tracks are shown, including start (green) and stop (red) codons with arrows indicating translation on the forward or reverse strand. *AtMα.2* overlaps with an uncharacterised protein (AT3G43686), predicted by NCBI to encode a pseudogene. Notably, *AtMα.2* and AT3G43686 are encoded on opposite strands, indicating that they do not share the same coding sequence, and supporting the classification of *AtMα.2* as a novel, previously unannotated MADS-box gene.

### A) *SlMα.5,* NC_015438.3: 69942216-69942839

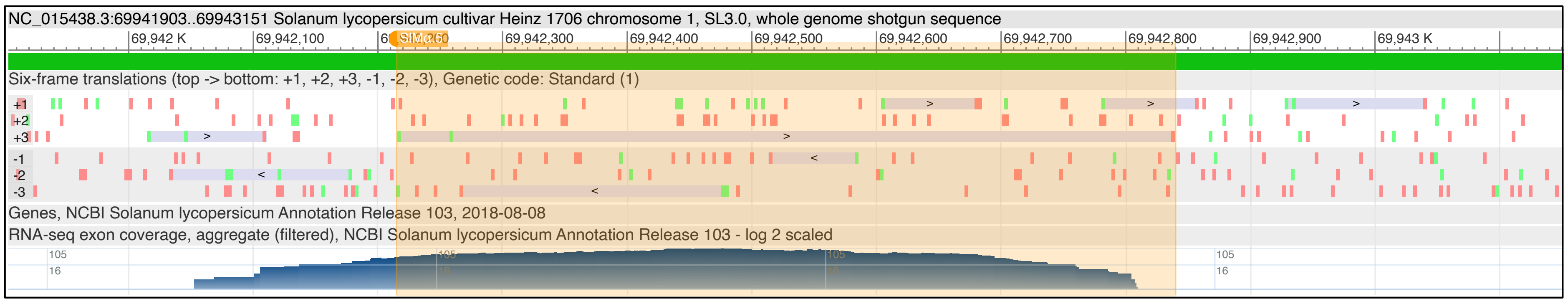


### B) *SlMα.4*, NC_015448.3:33097339-33097803

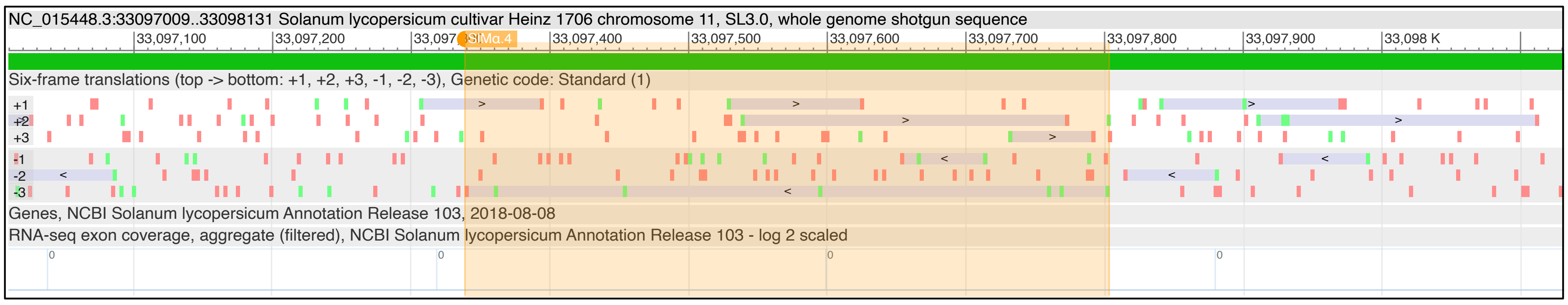


### C) *SlMα.1,* NC_015445.3:55191889-55192383

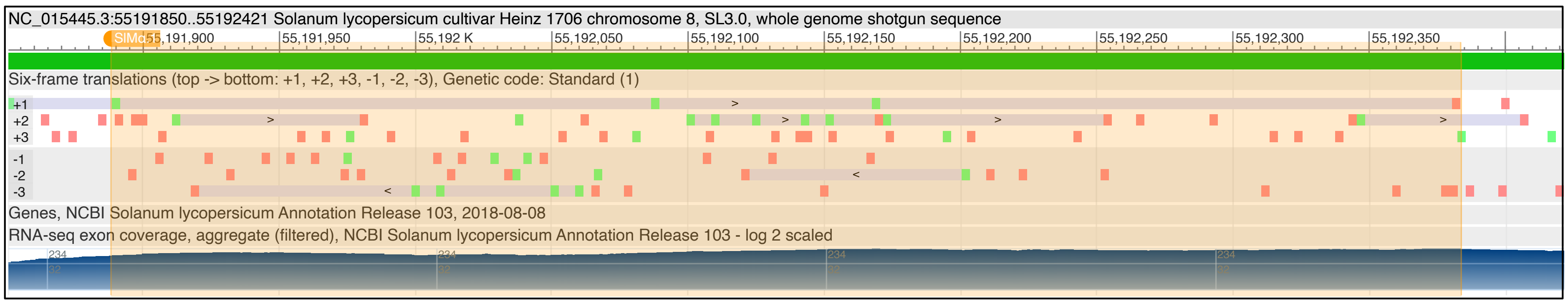


### D) *SlMα.2,* NC_015440.3: 69717841-69718374

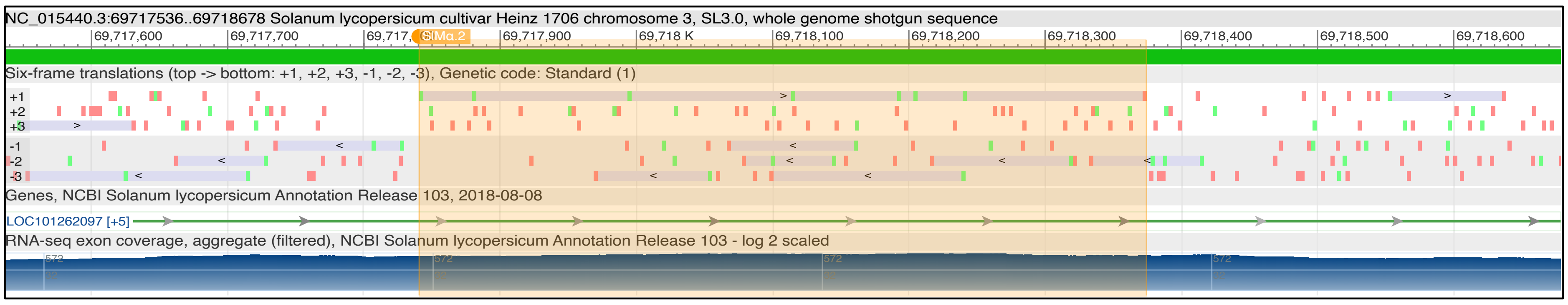


**Figure S5. Genomic position of newly predicted MADS-box genes in *S. lycopersicum* using GENE-FAM.** Novel predictions *SlMα.5* (A), *SlMα.4* (B), *SlMα.1* (C) and *SlMα.2* (D) are represented. Chromosomal locations are provided for each gene. Six-frame translation tracks are shown, including start (green) and stop (red) codons with arrows indicating translation on the forward or reverse strand. No previously annotated genes are found to overlap with any novel MADS-box gene prediction. Aggregated RNA-seq exon coverage tracks are shown. Overlapping RNA-seq coverage provides additional support for *SlMα.5*, indicating active transcription of the locus.

### A) *CsMα.1-C,* NC_044378.1:67860821-67861420

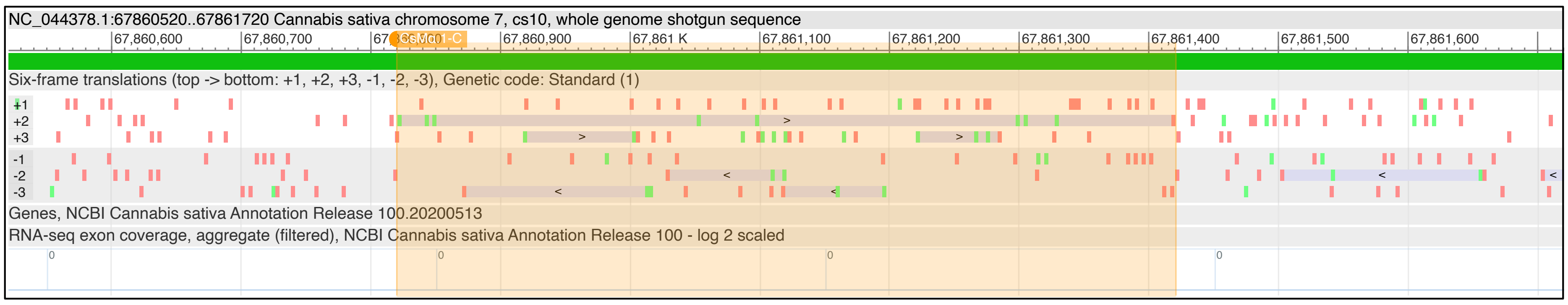


### B) *CsMα.32-C*, NC_044377.1:77156860-77158146

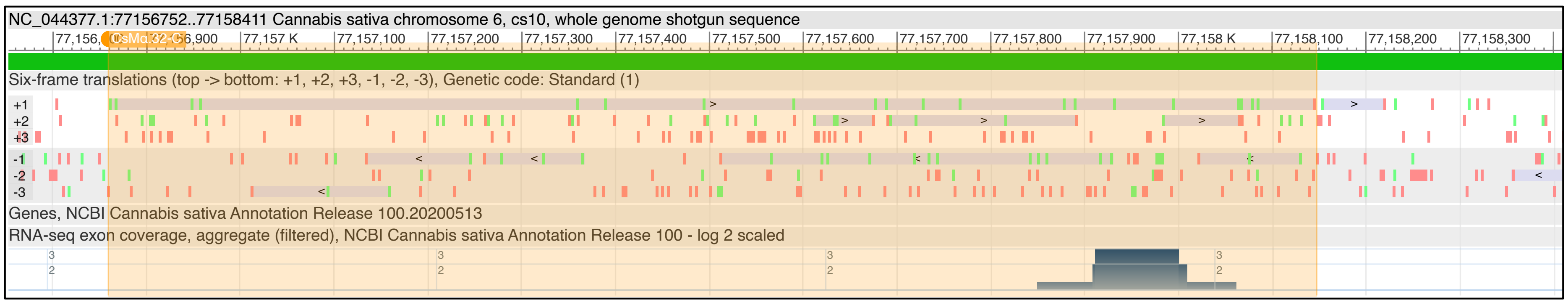


### C) *CsMα.22-C,* NC_044378.1:67881412-67882508

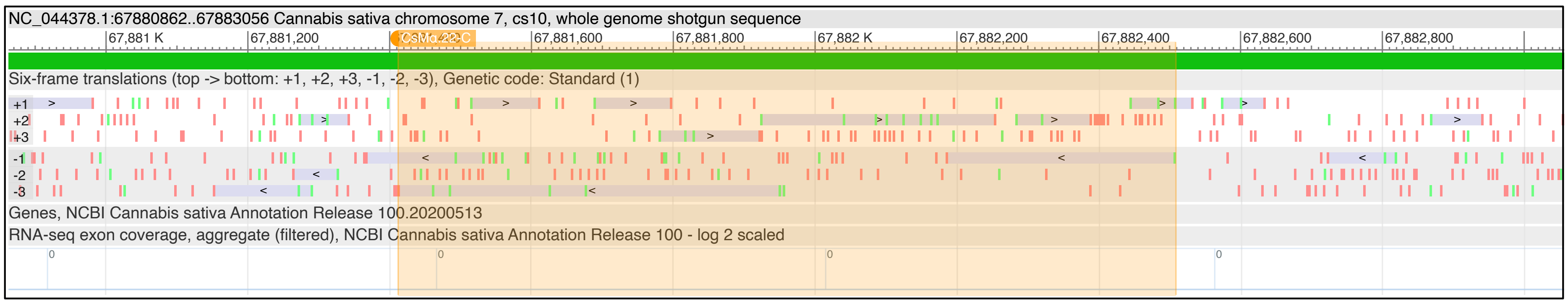


**Figure S6. Genomic position of newly predicted MADS-box genes in *C. sativa* using GENE-FAM.** Novel predictions *CsMα.1-C* (A), *CsMα.32-C* (B) and *CsMα.22-C* (C) are represented. Chromosomal locations are provided for each gene. Six-frame translation tracks are shown, including start (green) and stop (red) codons with arrows indicating translation on the forward or reverse strand. No previously annotated genes are found to overlap with any novel MADS-box gene prediction. Aggregated RNA-seq exon coverage tracks are shown, though support was minimal or absent for most loci.

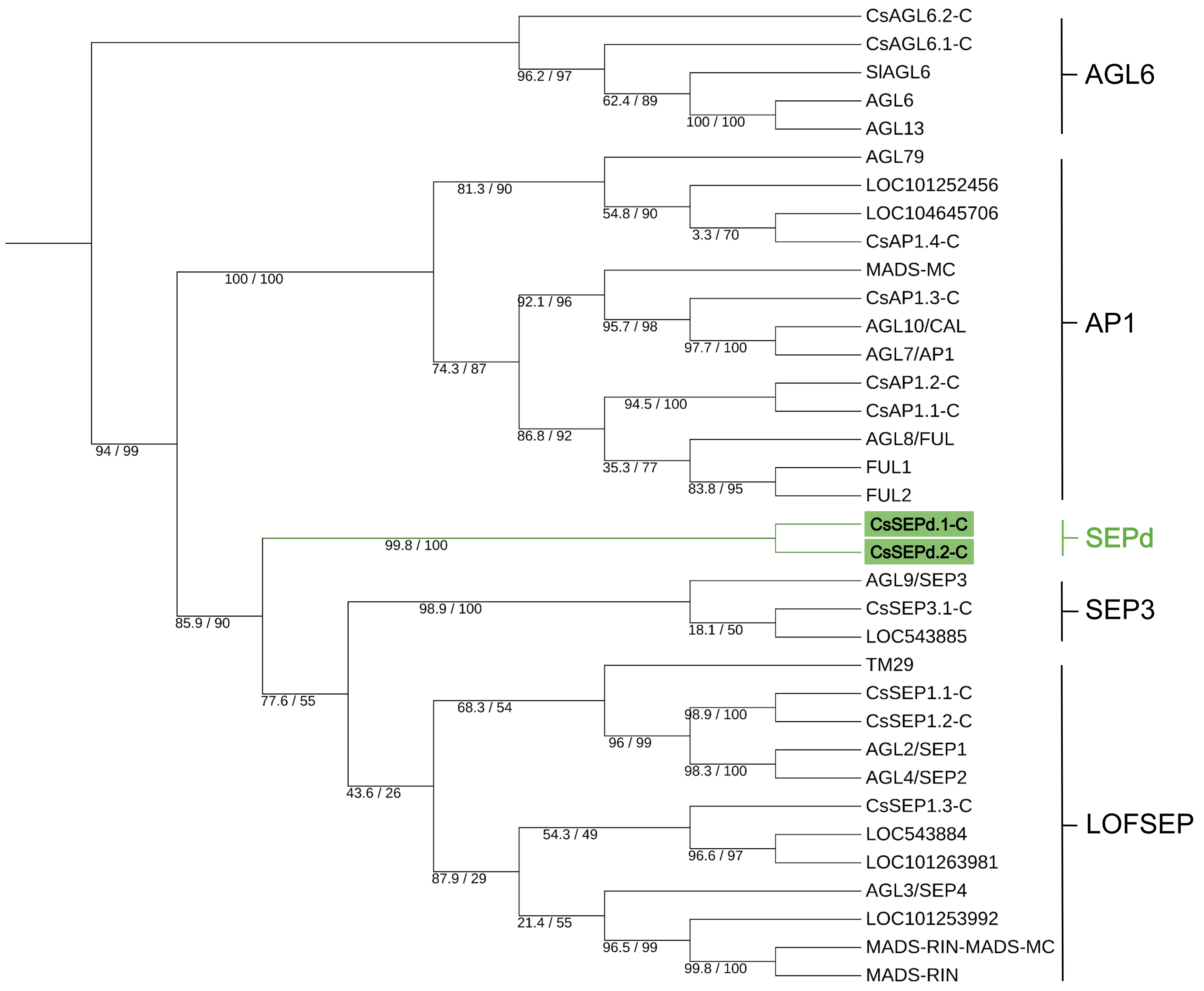


**Figure S7. Maximum likelihood phylogeny of proteins from the SEPd, SEP3, LOFSEP, AGL6, and AP1 subfamilies from *A. thaliana, S. lycopersicum*, and *C. sativa* (CBDRx cultivar).** High bootstrap values support the grouping of SEP-divergent (SEPd) proteins as a sister clade to the SEP3 and LOFSEP subfamilies.

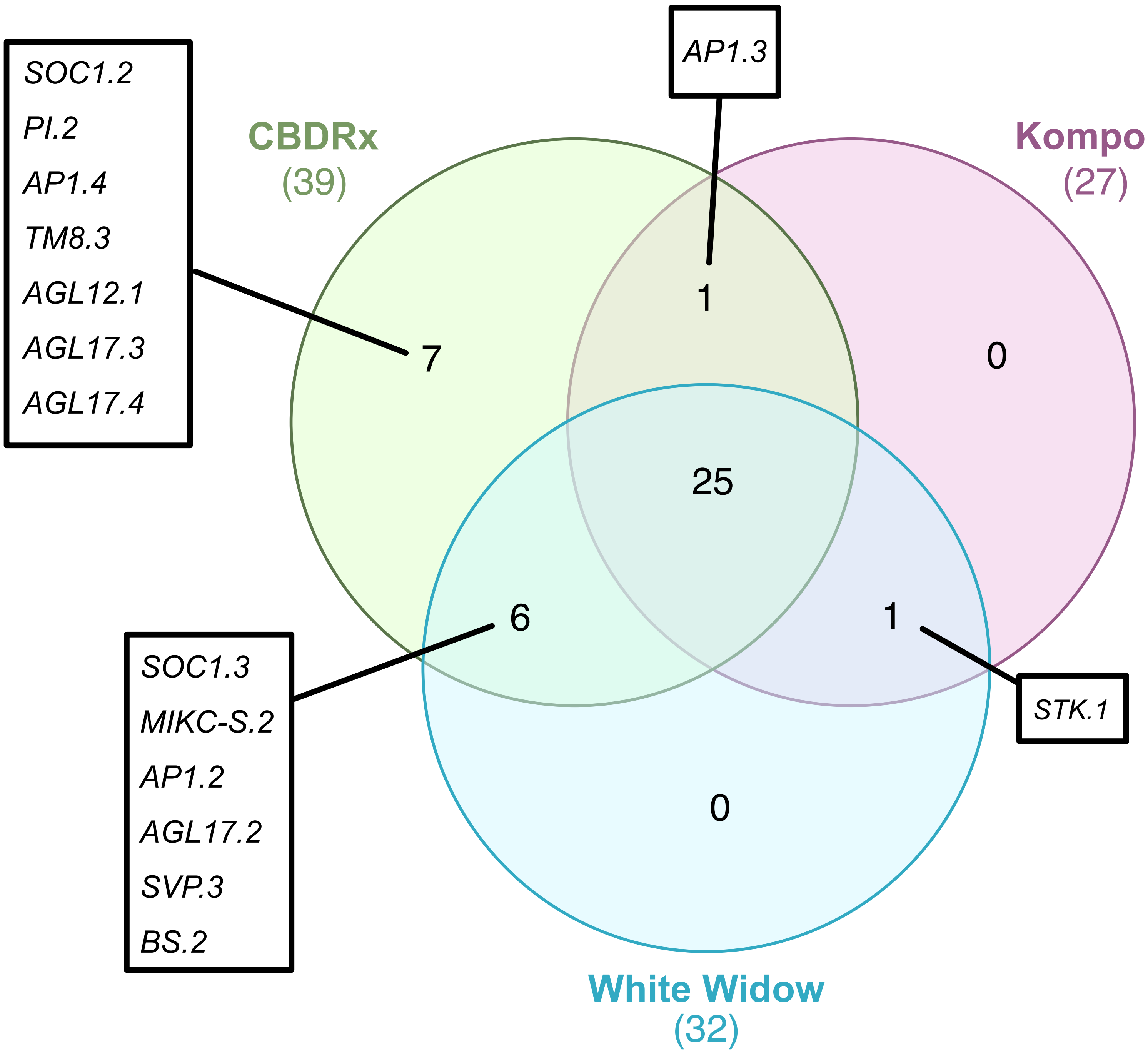


**Figure S8. Venn diagram illustrating conserved and cultivar specific Type II MADS-box gene repertoires across the CBDRx, 'White Widow' and 'Kompolti' *C. sativa* cultivars.** Haplotype specific gene sets for 'White Widow', and 'Kompolti' were combined to represent cultivar level repertoires.

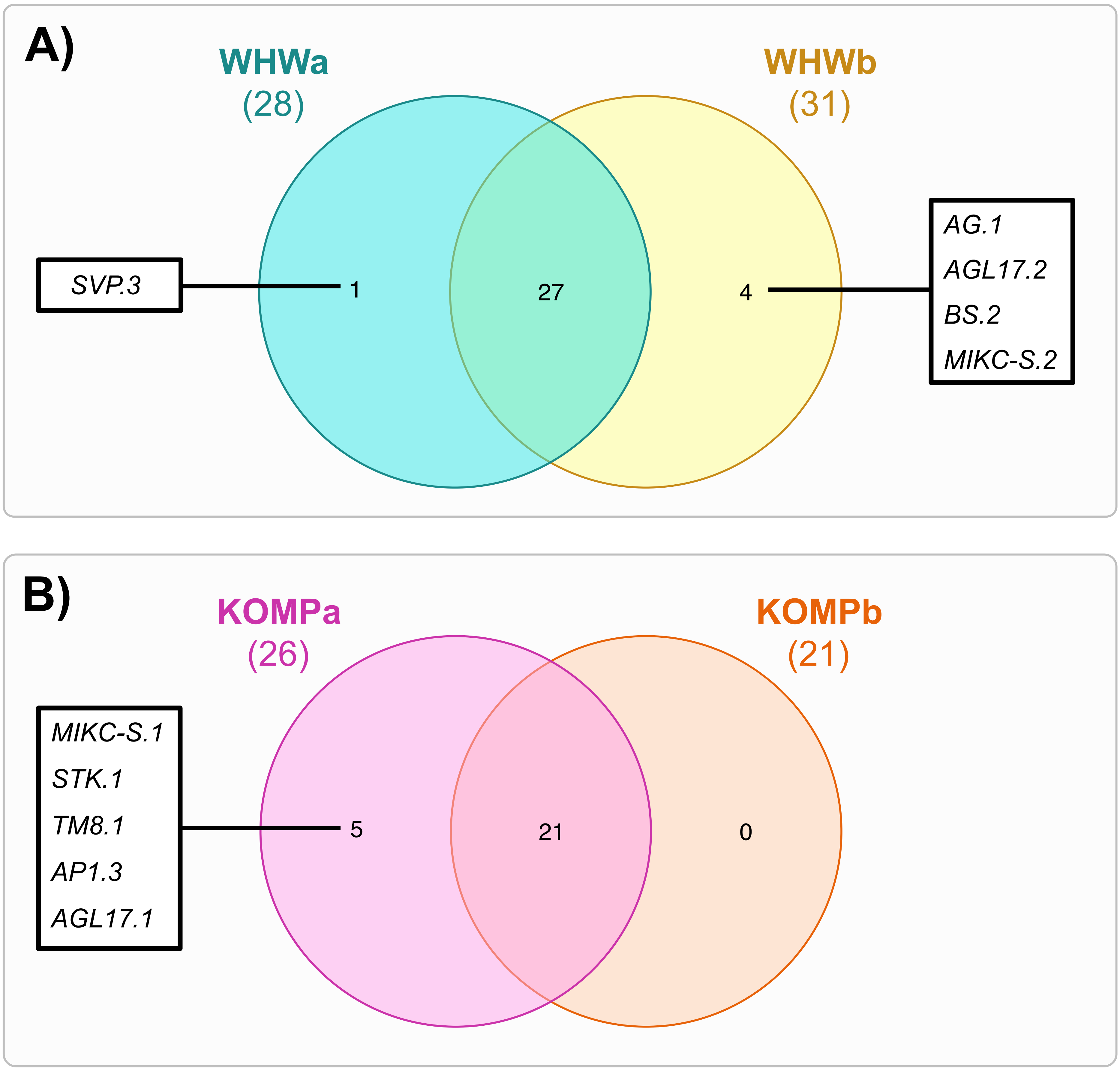


**Figure S9. Venn diagram illustrating conserved and haplotype specific Type II MADS-box gene repertoires across the (A) 'White Widow' and (B) 'Kompolti' cultivars.** Haplotypes are labelled as WHWa/WHWb and KOMPa/KOMPb, respectively.
